# Exploring mate choice and male strategies in a polymorphic moth

**DOI:** 10.1101/2024.08.20.608604

**Authors:** Eetu Selenius, Chiara De Pasqual, Matleena Hänninen, Liisa Kartano, Sandra Winters, Johanna Mappes

## Abstract

Color polymorphisms in natural populations often reflect the interplay between various selective pressures, such as natural and sexual selection. In this study, we investigate the dynamics of sexual selection operating on color polymorphism in wood tiger moths under different ecological contexts. Wood tiger moths exhibit polymorphism in male hindwing coloration, with individuals possessing one or two dominant W alleles displaying two forms of white coloration that differ in their UV reflectance (WW, Wy), while those with two recessive y alleles exhibit yellow coloration (yy). Females carry the color alleles, but do not express them phenotypically. We performed two mate choice experiments that simulated two ecological conditions: one with limited morph availability and low male encounter rates and the other with all morphs present and high potential for male encounters. We demonstrate that WW males consistently experience higher mating success compared to yy males, irrespective of the presence of Wy males and male encounter rates. Surprisingly, mating with a WW male does not confer direct reproductive benefits to females in terms of lifetime reproductive success; instead, Wy females exhibit overall higher reproductive success regardless of their mating partner. Although the precise mechanism driving the higher mating success of WW males remains unclear, a temporal decline in mating success of WW males suggests potential differences in male mating strategies. Our findings support the hypothesis that sexual selection contributes to the maintenance of polymorphism, revealing a stable mating advantage of a particular color morph that may be offset by other selective forces.

## Introduction

Understanding the mechanisms that can generate and maintain intrapopulation phenotypic variation is one of the key topics in evolutionary biology (Ford 1945, Barton and Turelli 1989). Natural selection is expected to erode variation in traits directly linked to fitness (Ford 1945, Lewontin 1974, Endler 1988), yet polymorphism in such traits is quite common in nature (Pruitt et al. 2008, Marchinko et al. 2014).

Polymorphism may be maintained through mechanisms such as frequency-dependent predation (Endler 1988, Punzalan et al. 2005), heterozygote advantage (Fisher 1923, De Pasqual et al. 2022) or sexual selection (Gadgil 1972, Wellenreuther et al. 2014). A lot of research on the role of sexual selection in maintaining polymorphism has focused on color polymorphisms, as they are relatively common across species (e.g., beetles (Osawa and Nishida 1992); frogs (Wang and Shaffer 2008, Rojas D et al. 2020); lizards (Sinervo and Lively 1996, Brock et al. 2020); moths (Hegna et al. 2015)). In addition, coloration has many functions linked directly or indirectly to fitness (warning coloration (Darwin 1859), mimicry (Poulton 1890), mating ability (Wilson et al. 1976), immunity (Cubaynes et al. 2022)) and is known to influence mate choice in several species (Meunier et al. 2011, Sefc et al. 2014).

Sexual selection can maintain color polymorphism in tandem with natural selection (Nokelainen et al. 2012), or in some cases on its own (Chunco et al. 2007), and the mechanism can vary from morph-specific assortative mating (several lizard species: Pérez i de Lanuza et al. 2013, Sacchi et al. 2018; Gouldian finch: Pryke 2009) to trade-offs in attractiveness and intersexual competition between color morphs (side-blotched lizards: Sinervo and Lively 1996; pygmy swordtail: Kingston et al. 2003) to morph-specific differences in breeding output (Eleonora’s falcon: Gangoso and Figuerola 2019). The multifaceted nature of sexual selection mechanisms results in a dynamic selective landscape that can vary spatially and temporally (Gosden and Svensson 2008). Therefore, the strength and direction of sexual selection on color morphs may fluctuate in response to ecological variation. For instance, changes in morph frequencies (Gossum et al. 2001, McLain 2005, Svensson et al. 2005, Gordon et al. 2015), population density (Cordero 1992), or the presence or absence of specific morphs (Corl et al. 2010) can alter the mating advantage of morphs. Such ecological influences can profoundly impact the maintenance or disappearance of multiple morphs within populations. However, the extent to which these ecological factors interact to influence sexual selection and color polymorphism remains poorly understood, highlighting a significant knowledge gap in our understanding of the evolutionary dynamics of these systems.

The wood tiger moth (*Arctia plantaginis*) provides an attractive model system to study the role of sexual selection in maintaining color polymorphism. Male hindwing coloration is determined by a single Mendelian locus with two alleles: the dominant W allele produces white coloration while the recessive y allele yellow coloration (Suomalainen 1938, Brien et al. 2022, Nokelainen et al. 2022) (Fig S1). While humans are not able to distinguish between homozygote white males (WW) and heterozygote males (Wy) as both are perceived as white, recent evidence suggests that conspecifics and key visual predators can separate WW from Wy moths based on differences in UV reflectance (Nokelainen et al. 2022). Females carry the same color alleles (i.e., W and y) but do not express them phenotypically, as their hindwing coloration varies between yellow and red (Nokelainen et al. 2022).

Thus far, studies have found differences between visible male color morphs (i.e., white vs yellow males) in terms of the effectiveness of the warning signal against visual predators (Nokelainen et al. 2012, 2014; Rojas et al. 2019; Winters et al. 2021), immune responses (Nokelainen et al. 2013), microbiome (Galarza et al. 2023) and mating success (Nokelainen et al. 2012; Gordon et al. 2015, 2018) where typically white males have a mating advantage over yellow males. However, the majority of studies have focused on differences between the visible color morphs without differentiating between the WW and Wy males. Yet very recent evidence of heterozygote advantage in terms of fertility, offspring survival and hatching success in Wy females (De Pasqual et al. 2022) highlights the importance of testing sexual selection at the genotype level.

Here, we set out to determine whether the three color genotypes (WW, Wy and yy) differ in their mating success and if mate choice is influenced by morph availability and male encounter rates. We approached these questions by measuring mating probability, mating behavior (i.e., latency to mate and female rejection rate) and reproductive output (fecundity, fertility, and hatching success) across male and female genotypes in two mate choice set-ups. Morph availability and the potential number of male encounters differed between the experiments, while sex ratio remained constant (1 ♀: 2 ♂). In the first experiment, we tested for differences between color genotypes in a low encounter rate scenario, where two genotyped males were offered simultaneously to a female in a direct pairwise comparison (WW vs Wy, WW vs yy, Wy vs yy). The second experiment took place in a large cage mating setting where 5 females and 10 males per genotype were present simultaneously to simulate a high encounter rate scenario. We expected overall that WW and Wy males would have a higher mating probability than yy males in both scenarios based on previous results (Nokelainen et al. 2012, Gordon et al. 2018). We also expected that heterozygote males and females would have higher fecundity and hatching success than the other color genotypes based on previous evidence of heterozygote advantage (Gordon et al. 2018, De Pasqual et al. 2022).

## Methods

### Study species

The wood tiger moth is a capital breeder species; adults do not feed, thus the resources accumulated at the larval stage are fundamental for individual development, reproduction, and survival (Tammaru and Haukioja 1996). In Finland, the species produces one generation per year with the mating season around June and July depending on the latitude. Females call for males by releasing sex pheromone which males perceive through their antennae (De Pasqual et al. *submitted*). After sensing the female pheromone, males cast the typical zigzag flight pattern to reach the female. As the male makes physical contact with the female, the female can express choice by either accepting a mating attempt or by rejecting the male by flapping her wings, moving away from the male or by dropping to the ground from the calling spot (personal observation).

### Stock maintenance

All individuals used in the experiments came from a laboratory stock that was established in 2013 at the Department of Biological and Environmental Science at University of Jyväskylä, Finland. Moths were reared in semi-natural conditions with natural lighting and controlled temperatures that matched outdoors temperatures (20 – 25 °C). New individuals are introduced to the stock from wild populations yearly. Within the stock, there are three genotype lines (WW, Wy, yy) that are constantly maintained for research purposes. Controlled matings are performed to maximize the genetic variability within genotype lines. In the laboratory setting, the wood tiger moth produces three generations per year and for these experiments we used moths from the second and third generation.

### Experimental settings

To measure the mating probability and reproductive fitness of the different color morphs, we performed mating trials in two experimental settings at the Department of Biological and Environmental Science (University of Jyväskylä). The experimental settings differed both in morph availability and male encounter rates.

#### a) Pairwise setting

To test for differences between male color genotypes under limited morph availability and low encounter rate, we performed a pairwise choice experiment. In each mating trial, two males in specific color genotype combinations were offered to a female (Table 1). These mating trials occurred between the months of June and August across four years; 2018 (n = 101), 2019 (n = 24), 2020 (n = 96) and 2021 (n = 53). Each year, the trials were done with moths from either the second (n = 154) or third (n = 120) annual laboratory generation of the stock. Out of the 274 mating trials performed, we retained 197 for the final analyses in which both female and male color genotypes were known based on the genotype lines (Table 1).

**Table 1.**
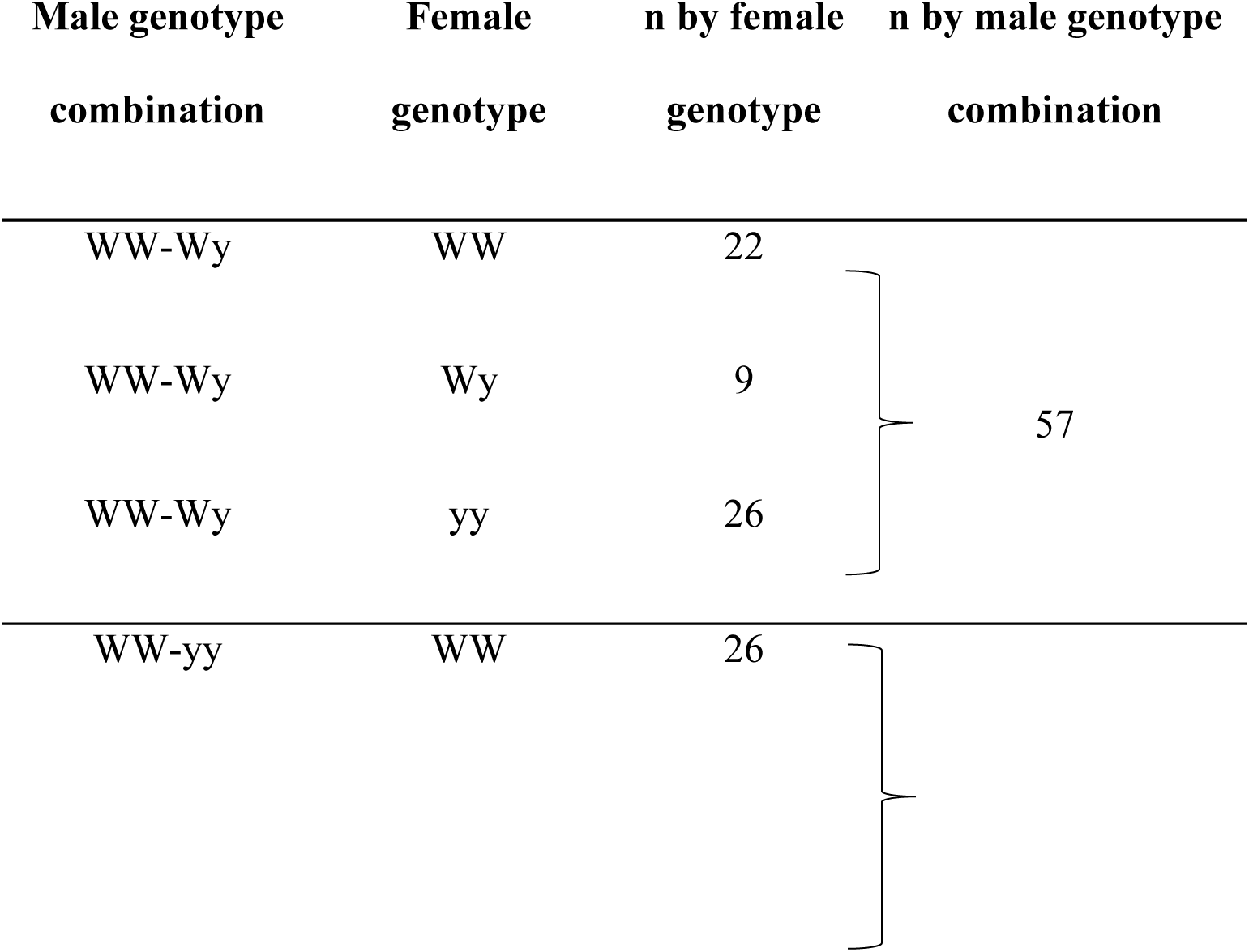
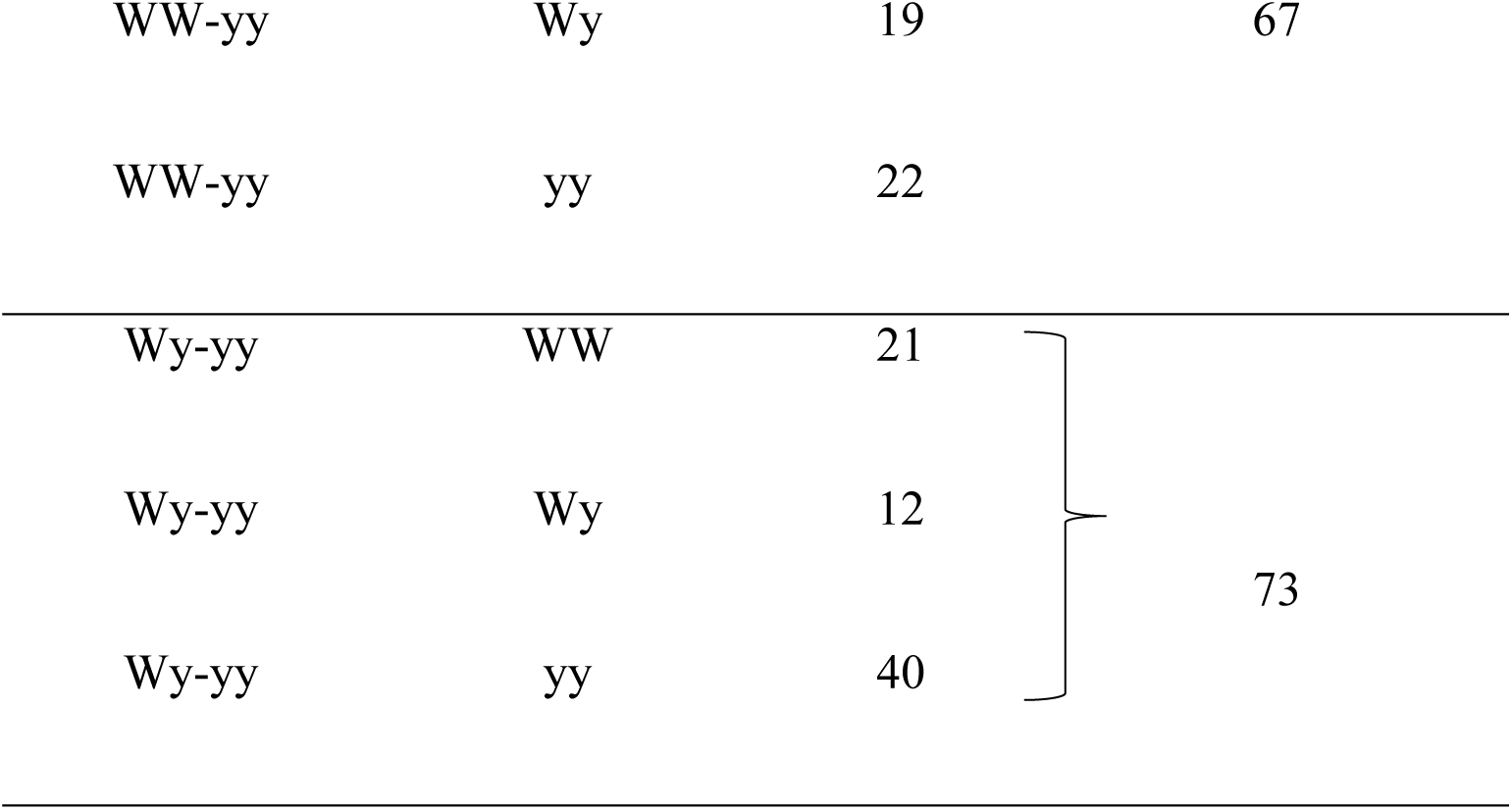
Final sample size of each pair combination tested in the pairwise choice experiment.

The trials were performed in plexiglass cages where the lid was replaced with regular glass to let UV into the experimental cage. According to a visual model, females are likely to differentiate between WW and Wy males based on UV reflectance (Nokelainen et al. 2022), which may therefore be important for female choice. In each cage (30 x 40 x 58 cm) we placed a twig from the bottom left corner to the top right corner, to offer an additional surface for the moths to rest on in addition to the walls and floor. We also placed a moistened sponge on the floor to offer water ad libitum. At 16:00h we released the moths into the cage and observed their behavior, mating probability, and mate choice until around midnight, when the moths become inactive as it becomes too dark for them to move. If a mating pair was formed during this time period, we allowed them to mate in the cage without disturbances for 15 minutes. Afterwards, the pair was transported from the experimental cage into a smaller plastic container (13 x 7 x 9 cm) to follow the reproductive fitness.

#### b) Large cage setting

To test whether mating probability and reproductive output differ across color genotypes under higher male encounter rates and with all morphs present at once, we performed mating trials where 15 females and 30 males were released into a large plastic cage (BILTEMA, 120 x 80 x 150 cm). In total, the experiment consisted of 15 trials and 675 individuals. The three color genotypes (WW, Wy and yy) were equally represented within each replicate, with five females and 10 males of each color morph. All individuals were identified using a four-dot marking system on the ventral side of their forewings and hindwings using permanent markers. Individuals were also marked according to their color genotype by adding a small dot of either blue, pink, or gold nail polish to their thorax in order to differentiate between WW and Wy males during observations. The color of the marking assigned to each color genotype was randomized between treatments. Individuals were marked during the day between 11:00h and 15:30h and released into the cage at 16:00h. The observational period lasted from 16:00h until the moths showed clear signs of decreased activity, typically around midnight (00:00h). Once a mating pair had formed in the cage, we allowed the pair to mate in the cage without disturbances for 15 minutes before moving the mated pair into a smaller plastic container (13 x 7 x 9 cm). In 2020, the experiment was conducted for only one night, but in 2021 the number of nights was increased to two to increase the number of matings.

### Variables considered

We measured the pupal weight of all individuals and used it as a proxy for adult weight. We also noted the age of each individual (in days) but did not include it in any of the models, as all adult moths were placed in a cold room (7 °C) on the same day they emerged from the pupa to halt the aging process, and they were selected for the experiments within one week of hatching (median 3 days).

For each experimental trial, we measured the following mating traits: A mating was considered successful if a mating pair had formed within the observation time and was assigned a value of 1. Otherwise, the mating was considered unsuccessful and assigned a value of 0. For each pair that successfully mated, we calculated the time in minutes it took for the pair to form (i.e., latency to mate). In addition, to test for potential assortative mating, we counted how many times each female color genotype mated with a specific male color genotype. As a proxy for female choice, we counted how many times a female rejected a male and how many times each male genotype was rejected.

After the mated pair separated, the male was removed from the plastic container while the female was left to lay eggs until the female died. To measure reproductive output, we counted the total number of hatched larvae as an estimate for lifetime reproductive success. The number of hatched larvae was counted 14 days after the first egg had hatched, as newly hatched larvae were too small to handle without the risk of causing them physical harm. Because of the high frequency of females who did not produce any living larvae, we also calculated the likelihood of producing viable offspring as a binary variable, with females assigned 1 if at least one of the laid eggs hatched and 0 if no eggs hatched. We also calculated the number of laid eggs as a proxy for fecundity. The overall hatching success was determined by dividing the number of larvae by the number of eggs.

### Statistical analysis

All statistical analyses were done using RStudio (version 2023.12.1, R version 4.1.2). We utilized Linear Models (LM), Generalized Linear Models (GLM) and Generalized Linear Mixed Models (GLMM) from package ‘*glmmTMB’* (Brooks et al. 2017). The overall effects of fixed effects were calculated using Wald Chi square tests implemented by ‘*Anova’* (package ‘*car’*: Fox and Weisberg 2019). For post-hoc pairwise comparisons between color genotypes, we used estimated marginal means (henceforth EMM, ‘*emmeans*’ function from package ‘*emmeans*’: Lenth 2021). Model distributions were evaluated using Kolmogorov-Smirnov tests and simulation-based dispersion tests provided by the package ‘*DHARMa’* (Hartig 2022). Plots were made in R using ‘*ggplot2’* (Wickham 2016).

We started by running full models with all relevant fixed effects, then generated reduced models using AIC for model selection, utilizing either the ‘*drop1’* or ‘*AIC*’ commands from base R. We kept either male or female color genotype always as the main effect, but included other fixed effects based on the model with the lowest AIC. For parts of the data where the sample size was smaller (according to Symonds and Moussalli 2010), we also calculated the corrected AIC values (AICc). If the difference in AIC was smaller than two for any models, we chose the simplest one according to Richards (2008). Full model selection process including all AIC values is in the supplementary material. All final models are listed in Table 2.

#### a) Weight

**Table 2.**
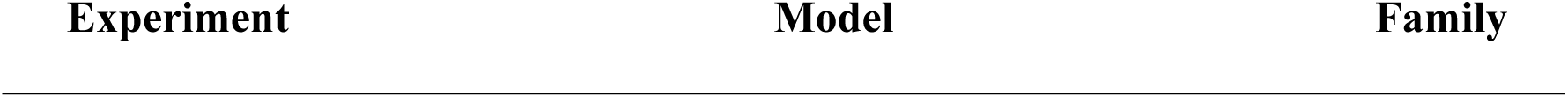
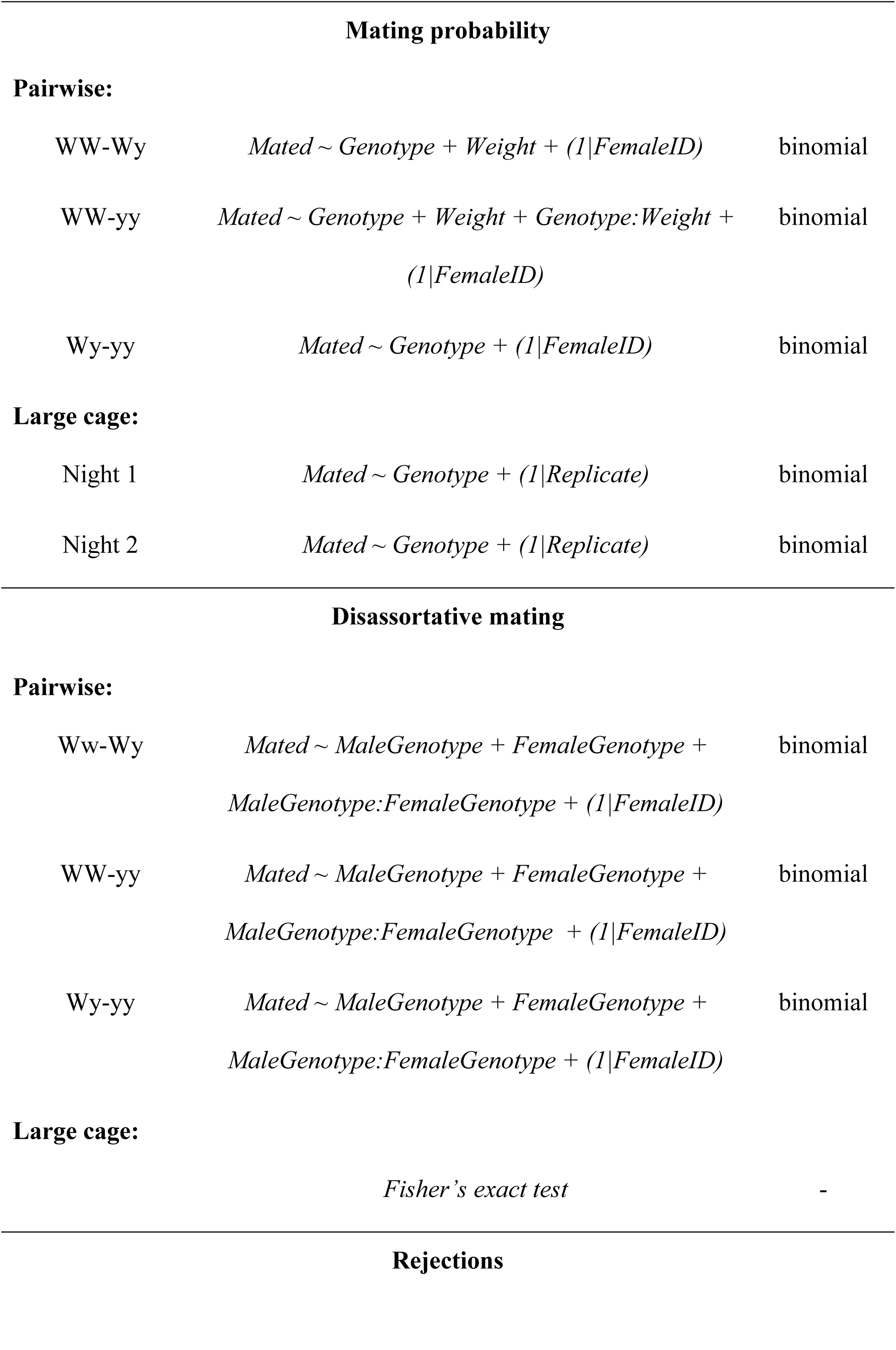
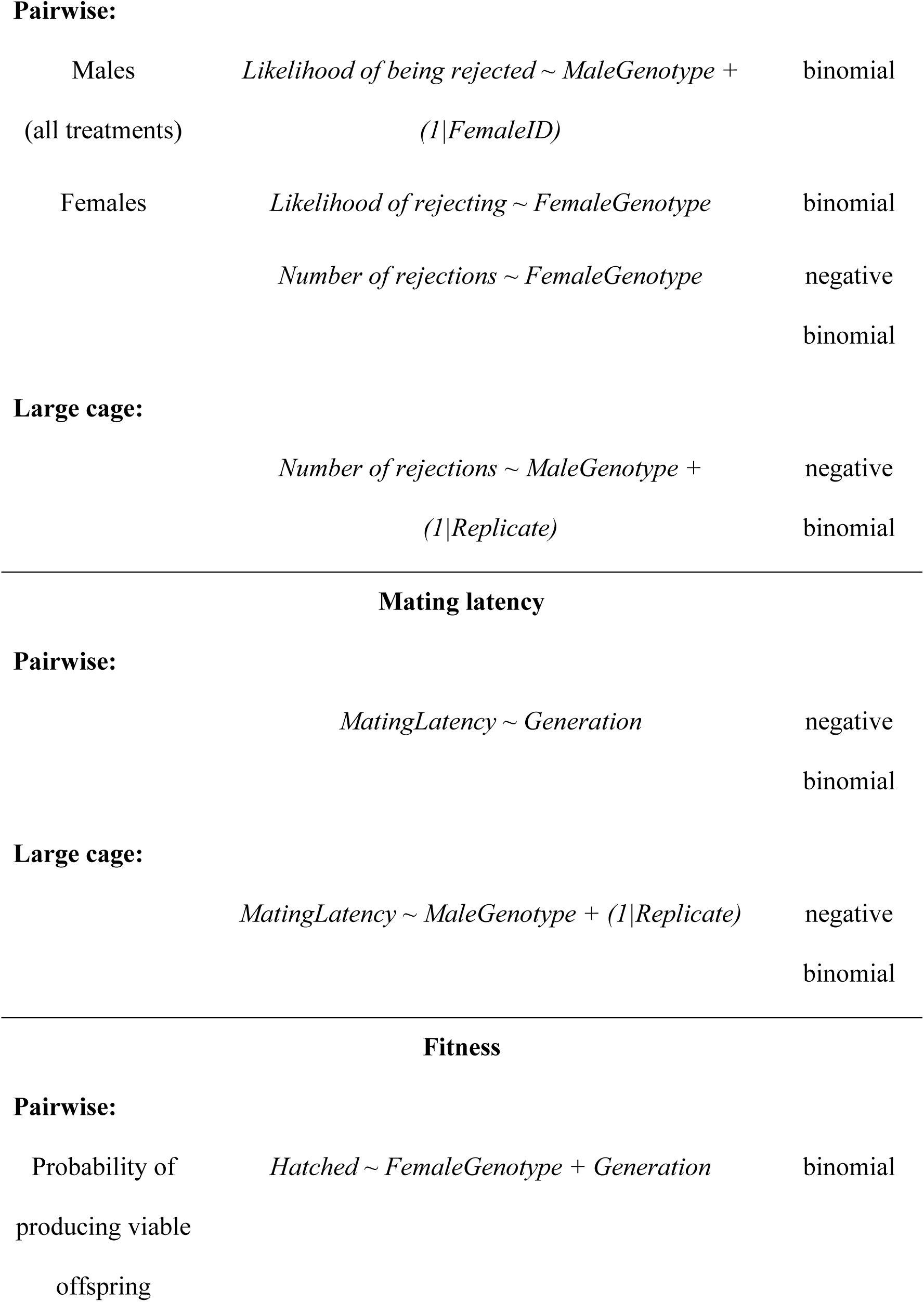
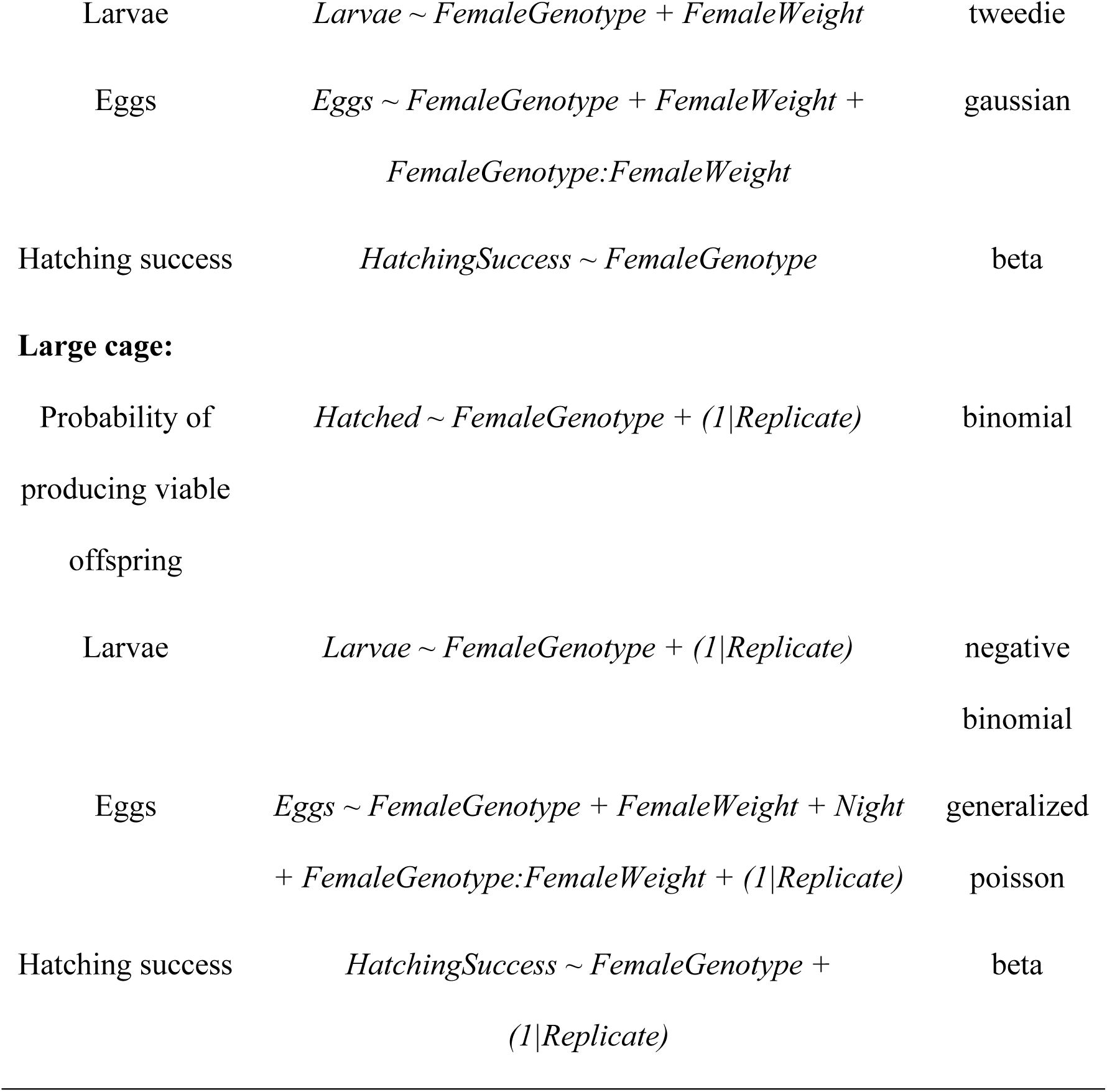
All statistical models that were selected using AIC. Response variable, fixed effects and random effects are listed under ‘Model’ section and the family distribution used in each GLM or GLMM is listed under ‘Family’.

| Mating probability |  |  |
| --- | --- | --- |
| **Pairwise:** |  |  |
| WW-Wy | $Mated \sim Genotype + Weight + (1/FemaleID)$ | binomial |
| WW-yy | $Mated \sim Genotype + Weight + Genotype:Weight + (1/FemaleID)$ | binomial |
| Wy-yy | $Mated \sim Genotype + (1/FemaleID)$ | binomial |
| **Large cage:** |  |  |
| Night 1 | $Mated \sim Genotype + (1/Replicate)$ | binomial |
| Night 2 | $Mated \sim Genotype + (1/Replicate)$ | binomial |
| Disassortative mating |  |  |
| **Pairwise:** |  |  |
| Ww-Wy | $Mated \sim MaleGenotype + FemaleGenotype + MaleGenotype:FemaleGenotype + (1/FemaleID)$ | binomial |
| WW-yy | $Mated \sim MaleGenotype + FemaleGenotype + MaleGenotype:FemaleGenotype + (1/FemaleID)$ | binomial |
| Wy-yy | $Mated \sim MaleGenotype + FemaleGenotype + MaleGenotype:FemaleGenotype + (1/FemaleID)$ | binomial |
| **Large cage:** |  |  |
|  | *Fisher's exact test* | - |
| Rejections |  |  |
| **Pairwise:** |  |  |
| Males | *Likelihood of being rejected ~ MaleGenotype +* | binomial |
| (all treatments) | *(1/FemaleID)* |  |
| Females | *Likelihood of rejecting ~ FemaleGenotype* | binomial |
|  | *Number of rejections ~ FemaleGenotype* | negative |
|  |  | binomial |
| **Large cage:** |  |  |
|  | *Number of rejections ~ MaleGenotype +* | negative |
|  | *(1/Replicate)* | binomial |
|  | *MatingLatency ~ Generation* | negative |
|  |  | binomial |
| **Large cage:** |  |  |
|  | *MatingLatency ~ MaleGenotype + (1/Replicate)* | negative |
|  |  | binomial |
| Probability of<br>producing viable<br>offspring | *Hatched ~ FemaleGenotype + Generation* | binomial |
| Larvae | $Larvae \sim FemaleGenotype + FemaleWeight$ | tweedie |
| Eggs | $Eggs \sim FemaleGenotype + FemaleWeight + FemaleGenotype:FemaleWeight$ | gaussian |
| Hatching success | $HatchingSuccess \sim FemaleGenotype$ | beta |
| **Large cage:** |  |  |
| Probability of<br>producing viable<br>offspring | $Hatched \sim FemaleGenotype + (1/Replicate)$ | binomial |
| Larvae | $Larvae \sim FemaleGenotype + (1/Replicate)$ | negative<br>binomial |
| Eggs | $Eggs \sim FemaleGenotype + FemaleWeight + Night + FemaleGenotype:FemaleWeight + (1/Replicate)$ | generalized<br>poisson |
| Hatching success | $HatchingSuccess \sim FemaleGenotype + (1/Replicate)$ | beta |

We standardized the weight (by centering the means and SD = 1) to include it in the interactions with discrete variables in the following analyses. We also tested for collinearity between color genotype and weight when both were included in the final statistical models (Supplementary material).

#### b) Male mating probability

To test for potential differences in male mating probability in the pairwise experiment, we fit three GLMMs, one for each treatment (WW-Wy; WW-yy; Wy-yy). We used mating probability as response variable modelled with a binomial distribution, and female ID as the random effect in all models. We used the male genotype, male weight, generation, and the interaction between male genotype and weight as initial fixed effects for model selection.

Since the trials in the large cage experiment lasted for one night in 2020 while they were carried out over two consecutive nights in 2021, we accounted for the effect of having an additional night of observations by running two sets of analyses: one using only the first night from both 2020 and 2021, and the other using only the second night from 2021. We tested for potential differences in the mating probability by fitting GLMMs with mating probability as response variable modelled with a binomial distribution and using the replicate as a random effect. Initially, we used male genotype, male weight and the interaction between male genotype and weight as fixed effects for model selection.

#### c) Disassortative mating

To test for potential disassortative mating in the pairwise set-up, we fit three GLMMs with binomial distributions, one for each treatment, with mating probability as response variable, the interaction ‘female genotype * male genotype’ as the main effect and female ID as a random effect.

For the large cage set-up, we tested for differences in the observed numbers of male genotypes that mated with each female genotype using a Fisher’s exact test.

#### d) Rejection events

For the pairwise experiment, we fit two GLMMs: one with the total number of rejections and the other the likelihood of rejection (0 or 1) as the response variable. We used male genotype, female genotype and their interaction as fixed effects and female ID as a random effect for model selection. For the males, we also fit a separate GLMM for each treatment with a likelihood of being rejected (0 or 1) as the binomial response variable, male genotype as the fixed effect and female ID as a random effect to test for differences between male genotypes within each treatment. Due to the low number of rejections, we could not include the interaction between male and female genotype in these models.

For the large cage experiment, we were unable to track rejections on an individual level, but we used a color genotype level summary of rejections (e.g., how many times WW females rejected yy males within each replicate). Using this number as the response variable, we fit a GLMM with female genotype, male genotype and their interaction as the fixed effects and replicate as a random effect. We then applied a pairwise post-hoc comparison using EMM.

#### e) Mating latency

For the pairwise set-up, we fit a GLM with mating latency as the response variable, and initially included male genotype, female genotype, their interaction, treatment, and generation as fixed effects.

For the large cage set-up, we fit an initial GLMM with mating latency as the response variable, male genotype, female genotype, their interaction and night as fixed effects, and replicate as the random effect.

#### f) Reproductive fitness

For both experiments, we fit four separate GLMs evaluating the likelihood of producing viable offspring, number of larvae, number of eggs and hatching success. For the pairwise experiment, we tested both female and male genotype, weight and generation as well as the interactions between female genotype and weight, and male genotype and weight as fixed effects, keeping female genotype as the main effect.

For the large cage experiment, we also used female and male genotype, weight and night as well as the interaction between female genotype and weight as fixed effects, keeping female genotype as the main effect.

## Results

### Male mating probability

In the pairwise set-up, male genotype had a significant effect on mating probability in the WW-yy treatment (χ^2^ = 7.351, df = 1, p = 0.007), with WW males having a higher mating probability than yy males (Fig 1a). The interaction between weight and genotype was also very close to significant (χ^2^ = 3.839, df = 1, p = 0.050), where an increase in weight increased mating probability of yy males but not WW males (EMM_WW-yy_: estimate = 1.95 ± 0.671, t = 2.903, p = 0.004).

**Figure 1.**
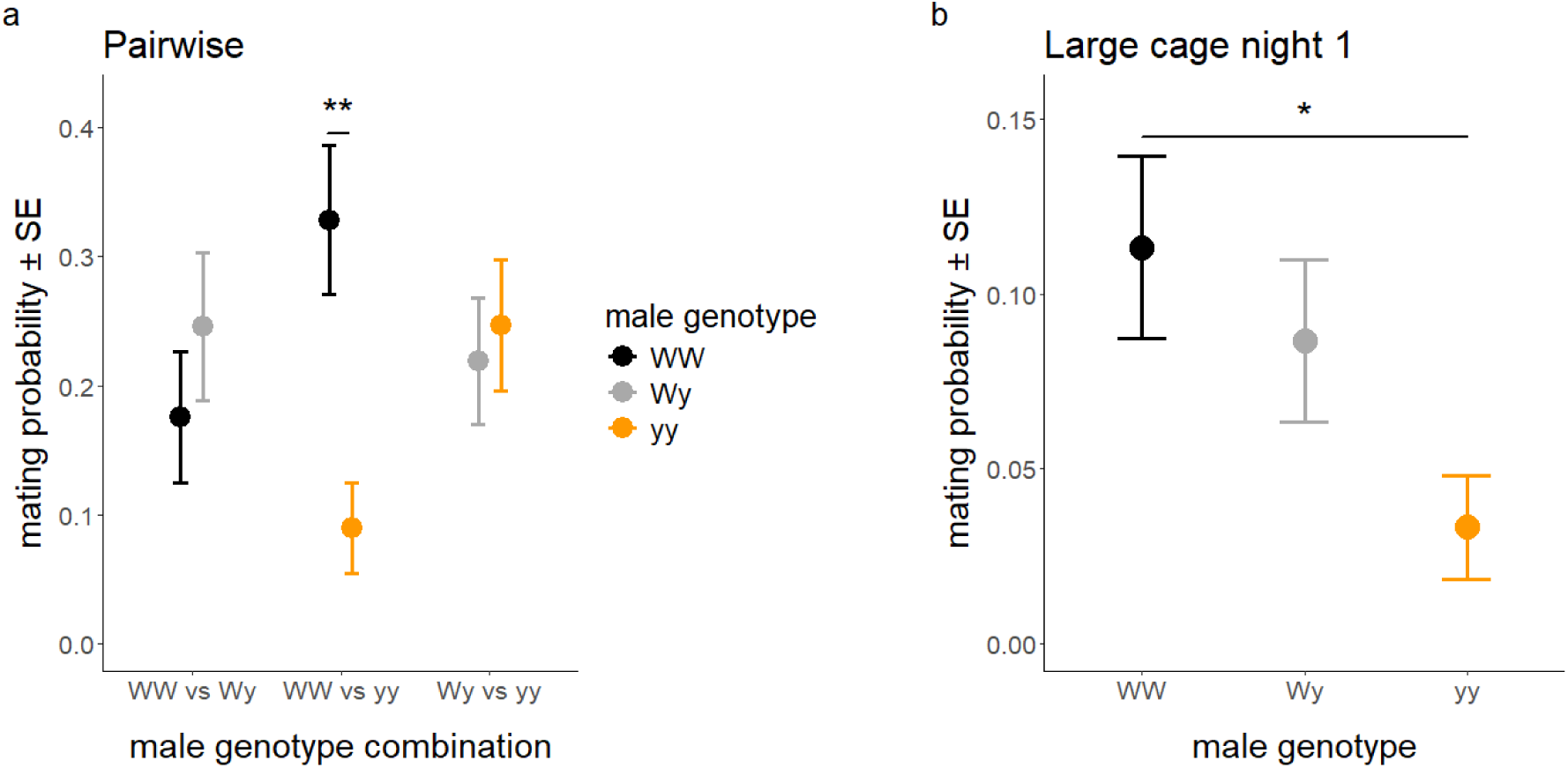
Mean male mating probability (± SE) a) within the different male genotype combinations in the pairwise set-up and b) between genotypes over the first night in the large cage set-up. a) WW males had a higher probability of mating when presented with a yy male whereas no differences were detected across the other genotype combinations. b) yy males were characterized by a lower probability of mating compared to WW males. Statistically significant differences are marked with asterisks: * = 0.05 < p < 0.01, ** = 0.01 < p < 0.001.

Male genotype did not have a significant effect on mating probability in the two other treatments, WW-Wy (χ^2^ = 0.772, df = 1, p = 0.380) and Wy-yy (χ^2^ = 0.090, df = 1, p = 0.764). Male weight significantly increased the mating probability of both genotype males in the WW-Wy treatment (χ^2^ = 4.431, df = 1, p = 0.035).

In the large cage set-up, when including only the first night of the experiment, male genotype had a significant effect on mating probability (χ^2^ = 6.367, df = 2, p = 0.041), with WW males having a higher mating probability than yy males (EMM_WW-yy_: estimate = 1.326 ± 0.526, t = 2.522, p = 0.032). Wy males also had a higher mating probability than yy males, although this difference was not significant (EMM_Wy-yy_: estimate = 1.023 ± 0.542, t = 1.886, p = 0.144) (Fig 1b).

Over the second night, male genotype also had a significant effect on mating success (). However, WW males had a significantly lower mating probability compared to Wy (EMM_WW-Wy_: estimate = −2.516 ± 1.060, t = −2.372, p = 0.049) and yy males (EMM_WW-Wy_: estimate = −2.568 ± 1.057, t = −2.430, p = 0.042) (Figure 2).

**Figure 2.**
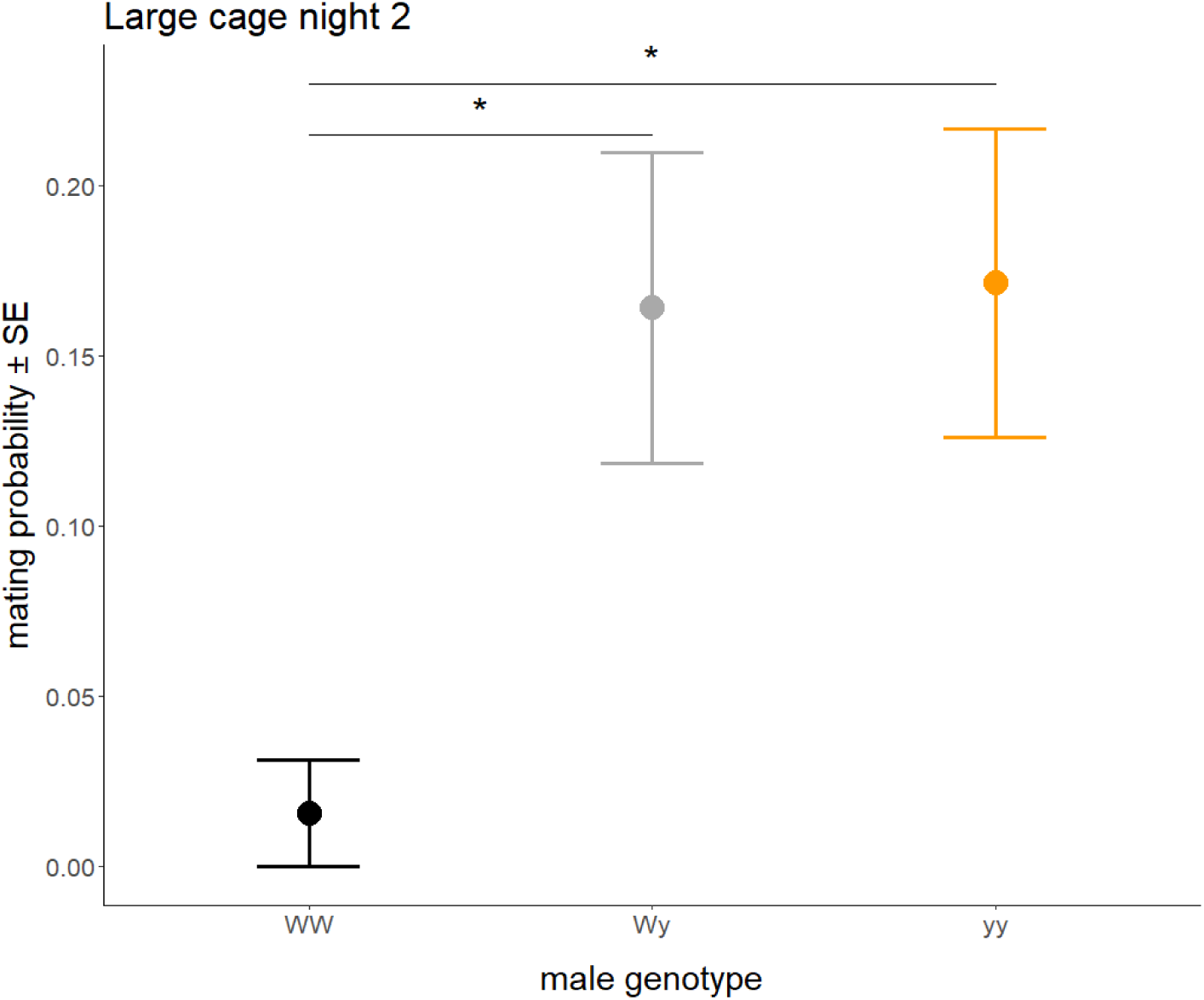
Mean male mating probability (± SE) between male color genotypes over the second night of the large cage experiment. WW males have a significantly lower mating probability than either Wy or yy males. Statistically significant differences are marked with asterisks.

### Disassortative mating

In the pairwise set-up, we found that WW females (EMM_WW-yy_: estimate = 1.727 ± 0.731, t = 2.362, p = 0.020) and Wy females (EMM_Wy-yy_: estimate = 21.100 ± 11500, t = 0.002, p = 0.999 justified by no Wy females choosing yy males) were more likely to mate with WW males when presented together with yy males, whereas they chose equally between the two males presented in the other male genotype combinations (EMM; all p > 0.05). yy females chose equally between males, showing no sign of mating preference (EMM; all p > 0.05) (Fig. 3). This suggests that WW and Wy females might have a stronger preference for WW male over yy males, whereas yy females do not present the same bias.

**Figure 3.**
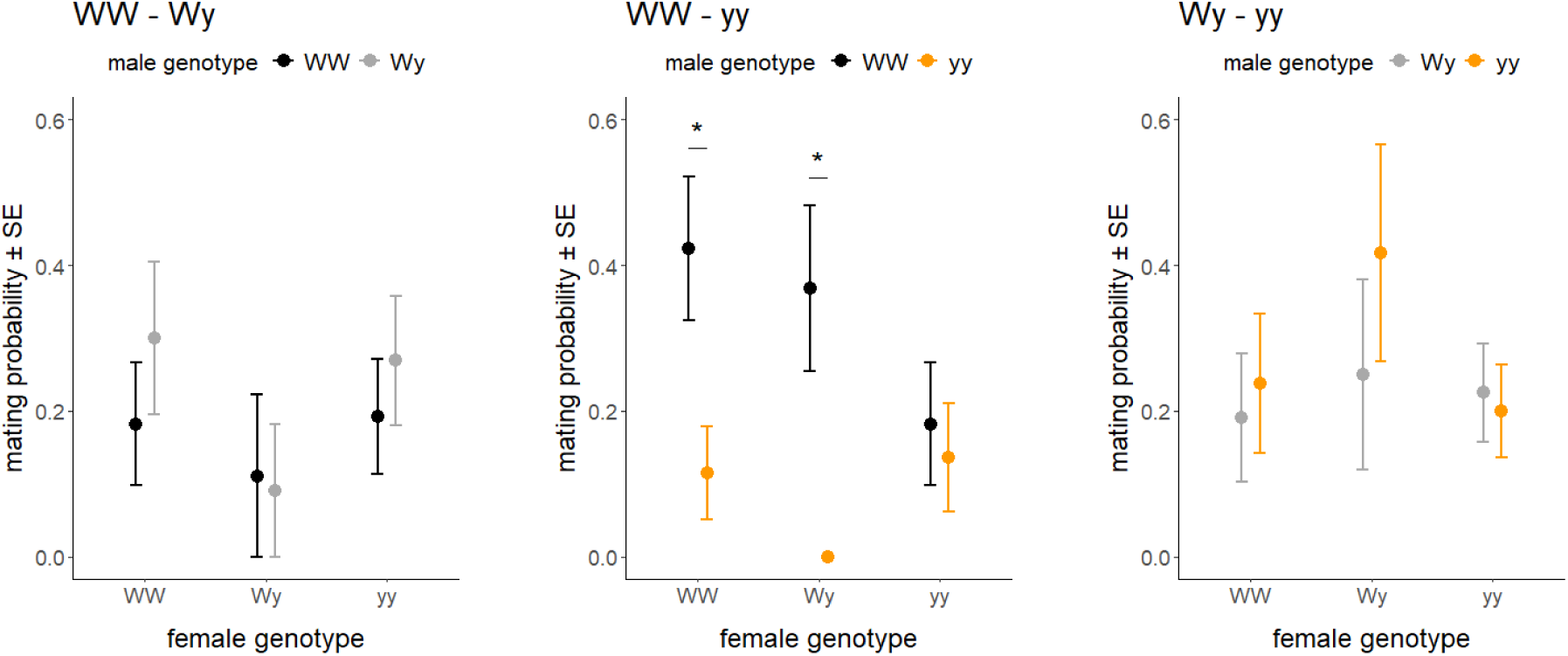
Mean male mating probability between male genotypes in different treatments, grouped by female genotype. Both WW and Wy females were more likely to mate with WW males over yy males when paired together (WW – yy), whereas yy females did not show a significant preference. Statistically significant differences are marked with asterisks.

In the large cage set-up, we did not find any differences between female genotypes in terms of which male genotype they were most likely to mate with (Fisher’s exact test; p = 0.713).

### Rejection events

In the pairwise set-up, we did not find a significant interaction between female and male genotype in either the total number (χ^2^ = 1.988, df = 4, p = 0.738) or the likelihood of rejections (χ^2^ = 2.218, df = 4, p = 0.696). Female genotypes also did not significantly differ in their likelihood of rejecting males (χ^2^ = 2.228, df = 2, p = 0.328), but there was a significant effect of genotype on the number of rejections (χ^2^ = 6.45, df = 2, p = 0.040), as yy females rejected on average less than WW females (GLMM_WW-yy_; estimate = −1.321 ± 0.555, z = −2.378, p = 0.017) (Fig. 4a).

**Figure 4.**
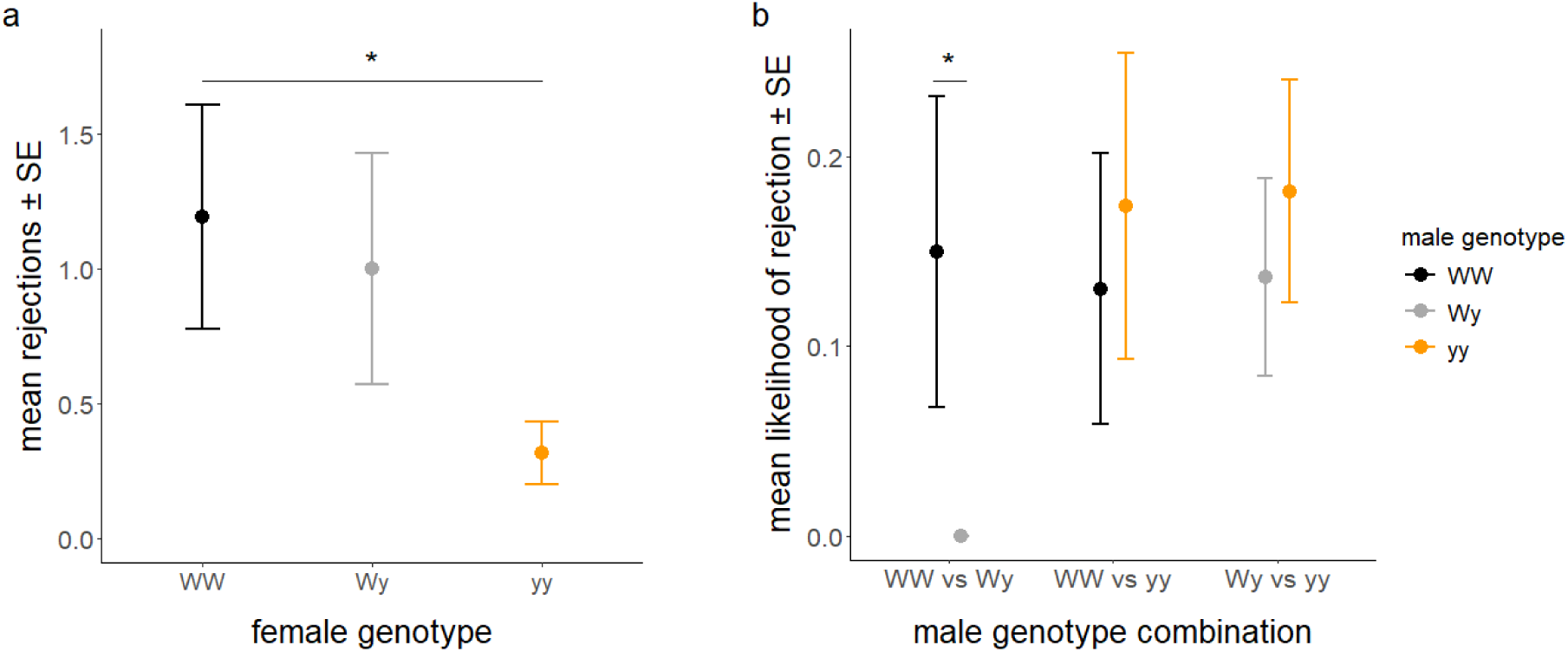
Average numbers of rejections among a) female genotypes, and b) male genotypes within each treatment in the pairwise experiment. Statistically significant differences are marked with asterisks.

WW males were significantly more likely to be rejected than Wy males when in direct competition (WW-Wy: χ^2^ = 0, df = 1, p = 1.000, justified by no Wy males being rejected). There were no significant differences between genotypes in the other two treatments (WW-yy: χ^2^ = 0.168, df = 1, p = 0.682; Wy-yy: χ^2^ = 0.338, df = 1, p = 0.561) (Fig 4b).

In the large cage set-up, we found no significant difference between how many times each male genotype was rejected (χ^2^ = 0.531, df = 2, p = 0.767).

### Mating latency

In the pairwise set-up, the mating latency was affected by the generation the mating took place (χ^2^ = 17.07, df = 1, p < 0.001). Third generation moths mated significantly faster (315 ± 12 minutes) than moths from the second generation (403 ± 12 minutes), implying an effect of seasonality.

Male color genotype had a significant effect on mating latency (χ^2^ = 15.892, df = 2, p < 0.001) in the large cage set-up. Despite having the highest mating probability, WW males mated significantly later then both Wy (EMM_WW-Wy_: estimate = 0.147 ± 0.037, t = 3.961, p < 0.001) and yy males (EMM_WW-yy_: estimate = 0.108 ± 0.042, t = 2.552, p = 0.036) during the observational period (Fig 5). Both Wy and yy males had overall fewer matings but mated earlier in the night than WW males.

**Figure 5.**
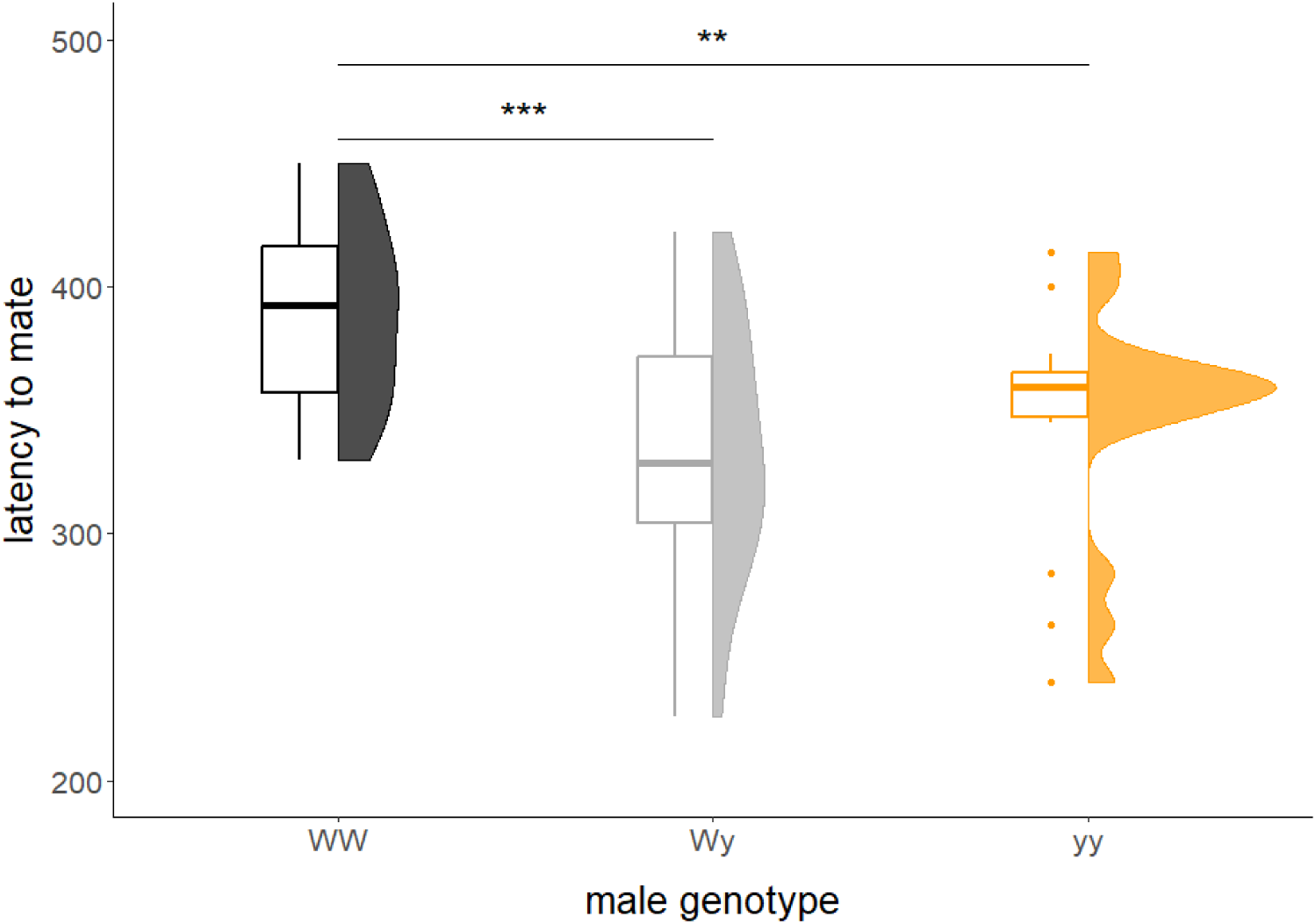
Mating latency of male genotypes in the large cage experiment. Statistically significant differences are marked with asterisks: ** = 0.01 < p < 0.001, *** = p < 0.001.

### Fitness

In the pairwise set-up, female genotype had a significant effect on the likelihood of producing any viable offspring (Fig 6a). yy females had a lower likelihood compared to Wy females (GLM; estimate = −1.813 ± 0.922, z = −1.967, p = 0.049). The likelihood of producing viable offspring also varied significantly between the two generations, being lower in the third generation (GLM; estimate = −2.251 ± 0.682, z = −3.302, p = 0.001). Female genotype did not have a significant effect on the number of larvae (χ^2^ = 2.751, df = 2, p = 0.253), but female weight had a significant positive correlation with the number of larvae (GLM; estimate = 0.277 ± 0.074, z = 3.72, p = 0.0002): heavier females produced more larvae (Fig 6b).

**Figure 6.**
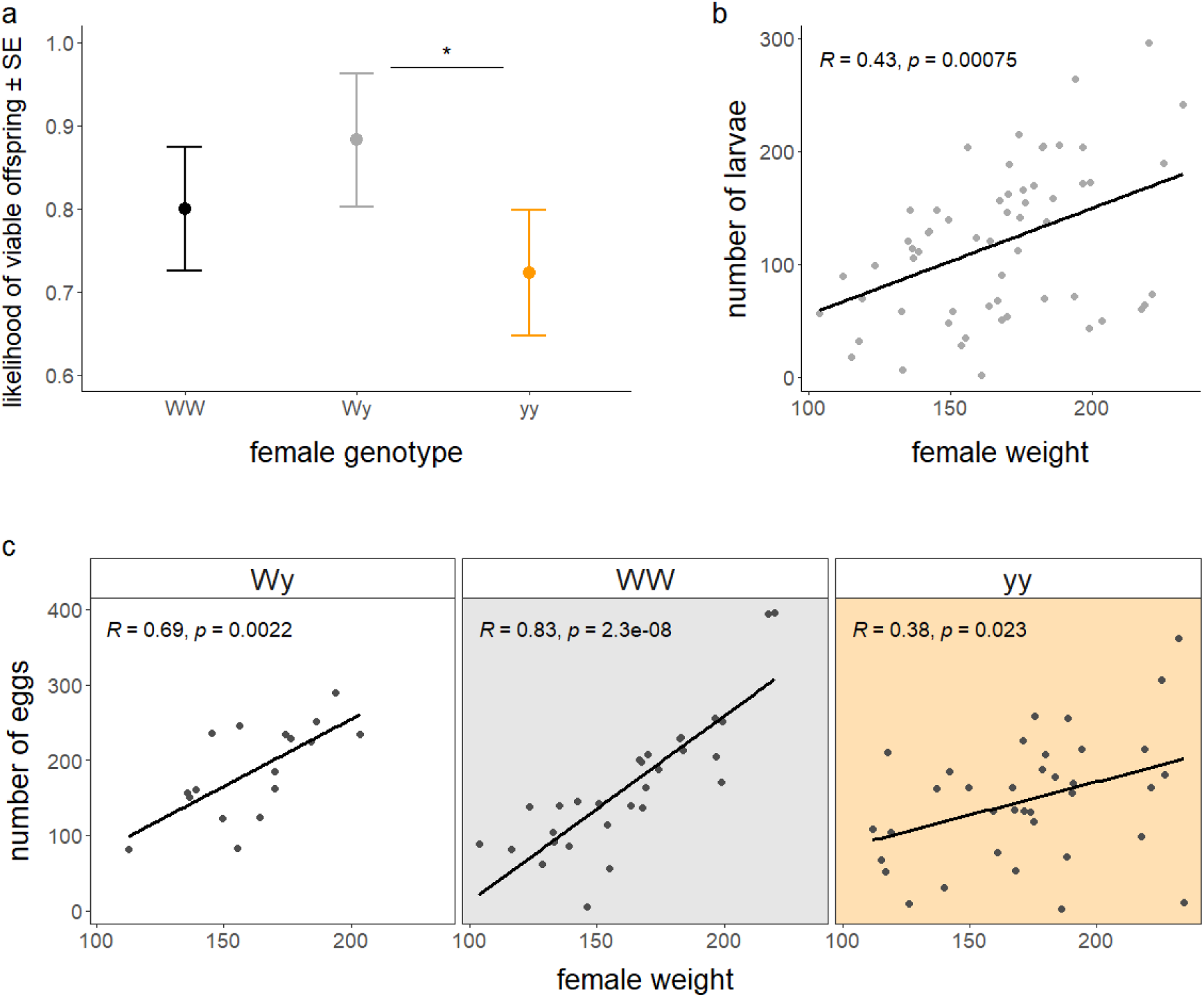
The effects of female genotype and weight on the reproductive output: effect of female genotype on the likelihood of producing viable offspring (a), effect of female weight on the number of larvae (b) and the interaction effect of genotype and weight on the number of eggs laid (c). Statistically significant differences are marked with asterisks.

The interaction between female genotype and weight had a significant effect on the number of eggs laid by females in the pairwise experiment (χ^2^ = 10.966, df = 2, p = 0.004). Weight had a stronger positive correlation with the number of eggs laid in WW females compared to yy females (GLM; estimate = 49.894 ± 15.223, z = 3.277, p = 0.001) (Fig 6c). Female genotype did not have a significant effect on the overall hatching success (χ^2^ = 1.474, df = 2, p = 0.479).

In the large cage experiment, female genotype did not have a significant effect on the likelihood of producing viable offspring (χ^2^ = 0.135, df = 2, p = 0.935) or on the number of offspring (χ^2^ = 0.328, df = 2, p = 0.849). The interaction between female genotype and weight had a significant effect on the number of eggs laid (χ^2^ = 8.859, df = 2, p = 0.012), as an increase in weight significantly increased the output of eggs in both Wy and yy females but not WW females (Fig 7). Night of the experiment also significantly affected the number of eggs laid, as females that mated on the first night of the experiment laid more eggs than females that mated on the second night (GLMM; estimate = −0.279 ± 0.111, z = −2.52, p = 0.012). Female genotype did not have a significant effect on the overall hatching success (χ^2^ = 2.432, df = 2, p = 0.296).

**Figure 7.**
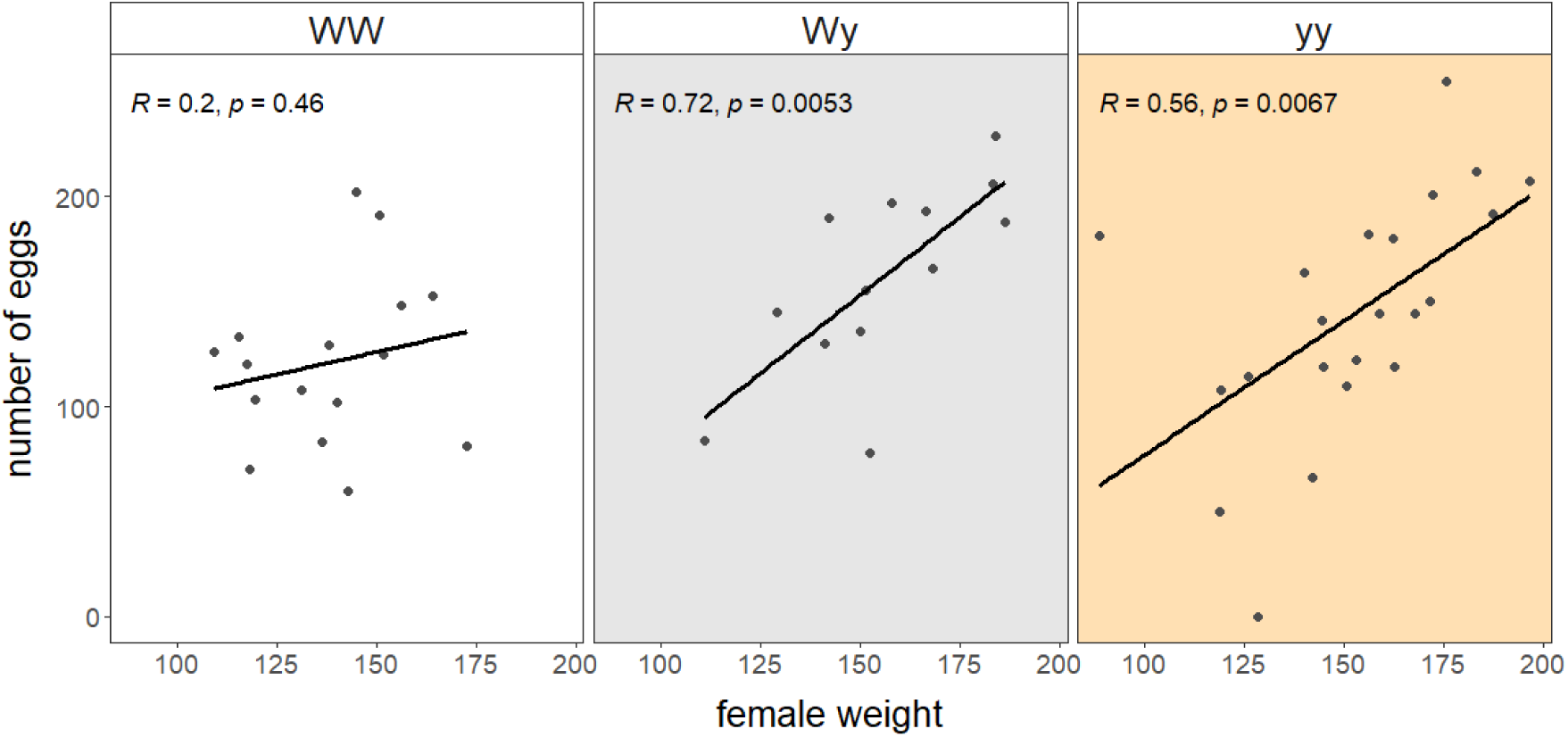
The interaction effect of weight and female color genotype on the number of eggs laid by females.

## Discussion

Sexual selection is known to contribute to the maintenance of color polymorphism in many species through several mechanisms of selection (*see* Wellenreuther et al. 2014).

Color morphs may experience balancing selection through trade-offs between natural and sexual selection (Nokelainen et al. 2012) or be maintained through negative frequency-dependent sexual selection (Svensson et al. 2009, Estévez et al. 2020) or other scenarios where the morphs’ mating success is context-dependent (Kvalnes et al. 2022). Better understanding how sexual selection operates on color polymorphic species under varying ecological contexts, may help us not only better understand how polymorphism is maintained, but also explain differences in color morph frequencies between different populations (Corl et al. 2010). Here, we provide evidence for a relatively stable morph-specific mating advantage in the wood tiger moth and discuss its role in maintaining color polymorphism.

We studied how polymorphic male wood tiger moths differ in their mating success in two different mating scenarios: one with limited morph availability and low male encounter rates (pairwise), and the other with all color morphs present and with higher potential male encounter rates (large cage). WW males had a significantly higher mating probability than yy males in both settings, although WW males’ mating probability became significantly lower than both Wy and yy males’ on the second night of the large cage experiment when the moths were allowed to mate for two nights in a row. WW males also had a significantly higher mating latency compared to the other two morphs in the large cage. In the pairwise experiment, only WW and Wy females preferred WW males over yy males, while all morphs had an equal probability of mating with a yy female. We saw no clear indications of active female mate choice, as all male color morphs were equally likely to get rejected by females. Finally, we showed that mating with a WW male did not give females a fitness advantage through enhanced reproductive success, as number of offspring (larvae) was not affected by male color morph: even though females that mated with WW males produced more eggs in the large cage, it was not translated to the number of offspring. However, females carrying both color alleles (Wy) had a significantly higher likelihood of producing viable offspring than yy females, indicating a form of heterozygote advantage (De Pasqual et al. 2022).

At first, the constant higher mating probability of homozygote white males may seem puzzling in terms of maintaining color polymorphism, as one could expect quick fixation of the W allele in populations. However, previous studies offer several explanations for how polymorphism could be maintained in the species: First, the mating advantage of WW males could be balanced by natural selection favoring the yellow (yy) morph (Nokelainen et al. 2012). This balancing selection could also explain some variation in morph frequencies across populations (Hegna et al. 2015), as the Scottish population associated with higher predation rates is monomorphic yellow (Nokelainen et al. 2013). Second, sexual selection could maintain polymorphism on its own if the WW advantage is driven by either assortative or disassortative mate choice (*see* Pérez i de Lanuza et al. 2013, Hedrick et al. 2016, Sacchi et al. 2018) or fluctuates based on morph availability (Sinervo and Lively 1996). We did not find convincing evidence of either assortative or disassortative mating, although both mating probability and rejections of male morphs varied slightly between the females carrying different color alleles in the pairwise set-up, indicating potential variation in female preference across color morphs. We also did not find evidence of mate choice fluctuating based on morph availability, which would be expected if male color polymorphism is maintained through non-transitive fitness advantages of the morphs, similar to the side-blotched lizard (Sinervo and Lively 1996). A WW mating advantage over yy males remained in both experimental settings, independent of the presence of Wy males. However, this advantage disappeared when the pair was not in direct competition, as both morphs had a relatively similar mating probability when paired with a Wy male. This suggests that one or more traits important to achieve copulation strongly differ between WW and yy males, and this difference stands out when these two male morphs are in direct competition and allows WW males to outcompete yy males.

As coloration is often genetically correlated with other fitness-related traits (McKinnon and Pierotti 2010), color morphs can differ from each other in multiple traits under sexual selection (e.g., alternative mating strategies: Sinervo and Lively 1996; breeding success: Gangoso and Figuerola 2019). We discuss here three possible traits whose variation between the male morphs may affect sexual selection in non-mutually exclusive ways. In Lepidoptera, mate choice can be based on a visual signal, a chemical signal or their combination (Phelan and Baker 1986, Iyengar et al. 2001, Robertson and Monteiro 2005, Costanzo and Monteiro 2007). According to color vision models, wood tiger moth females can discriminate between all male color genotypes based on different UV reflectance of WW and Wy males (Henze et al. 2018, Nokelainen et al. 2022). It is thus possible that, when in direct comparison, specific chromatic and luminance differences of WW and yy males interact to skew female choice towards WW males.

Female choice can also be based on male chemical signals, such as short-range sex pheromones, typically emitted by a male once in close proximity to a female (Birch et al. 1990; Iyengar et al. 2001). Variation in the pheromone blend emitted by males can be associated with the reproductive potential of the male (Phelan and Baker 1986, Iyengar et al. 2001), individual identity (Nieberding et al. 2012) or male size (Phelan and Baker 1986, Iyengar et al. 2001) and can affect female choice. Preliminary chemical analyses in wood tiger moth males found pyrrolizidine alkaloids (i.e., PAs, the metabolic precursors of hydroxydanaidal (Conner et al. 1981, Schulz et al. 1993)) in the legs (Winters et al. *in prep.*), which together with behavioral observations of males rubbing their legs on females during courtship (ES and CDP Pers. Obs.) hint to a potential involvement of male short-range sex pheromone in the mating process. Future investigations are needed to test whether potential associations between male sex pheromones and color alleles affect between-individual interactions, as we have recently shown for female sex pheromones (De Pasqual et al. *submitted*). Finally, the greater mating success of WW males over yy males may be linked to behavioral differences.

Many species are characterized by variance in sexual behavior, or the existence of distinct alternative mating strategies (Sinervo and Lively 1996, Kingston et al. 2003, Shuster and Wade 2003, Pryke and Griffith 2006). Our results indicate potential behavioral differences between male color morphs in terms of mating latency and temporal differences in mating probability, especially in the large cage experiment.

Despite their overall higher mating probability during the experiment, WW males had significantly higher latency to mate compared to both Wy and yy males. Also, when the experiment was continued for two nights in a row in 2021, we saw a complete turnover in morph-specific mating probability, as WW males rarely mated on the second night. WW males may display a mating strategy where they gain advantage by using energy early and outcompeting other males on the first night females start calling but losing that advantage over time. Lower mating latency of yy males may be explained partly by their apparent ability to locate females faster than the other morphs, as was shown by De Pasqual et al. (submitted). This latency effect may not be visible in the pairwise experiment due to the smaller size of the mating boxes, which could weaken the importance of mate location, or due to it being masked by the strong effect of generation on mating latency.

Sexual selection usually promotes mating with individuals of higher fitness, often selected via male-male competition or female choice (Darwin, 1859). We approximated the level of female choosiness by measuring the number and likelihood of rejections by females and found that both WW and Wy females rejected more than yy females in the pairwise experiment. The lower rejection rate of yy females may partly explain why we found no difference in their likelihood to mate with either WW or yy males. Since yy females are characterized by overall lower mating probability and reproductive success (De Pasqual et al. 2022), it is possible they are less choosy to maximize their mating chances. Female choosiness is expected to be lower when mating is not guaranteed (Kokko and Mappes 2005), and it may indeed be costly for yy females to reject a male, especially when the likelihood of encountering further males is low in the pairwise experiment.

Quantifying the reproductive success of color morphs is crucial to understanding the full role of sexual selection, as high mating probability may not be correlated with high reproductive output due to trade-offs between pre- and postcopulatory investments (Simmons et al. 2006, Durrant et al. 2016). Considering the reproductive output of both males and females also allows us to disentangle which sex plays a larger role in reproductive fitness. Although WW males were more likely to form a mating pair (especially compared to yy males), we did not find that females that mated with WW males accrued short-term fitness benefits, at least in terms of fertility and hatching success. However, as also discussed by Palokangas et al. (1992), there remains the possibility that offspring of preferred males have higher viability or mating probability. While our results showing a lack of direct and indirect benefits are in line with a previous experiment with the wood tiger moth (Santostefano et al. 2018), neither study tracked offspring fitness in the next generation. Further investigation on the offspring viability or mating probability is therefore needed before completely ruling out the possibility of indirect fitness benefits in this species.

While we did not find a clear contribution of male color genotype to reproductive fitness, Wy females had a higher probability of producing viable offspring compared to yy females, when mating in the pairwise set-up. This finding is partially in line with our prediction of Wy females having higher reproductive fitness, as found in De Pasqual et al. (2022). The lack of a comparable heterozygote advantage effect in the large cage set-up may be due to the smaller sample size, or as discussed in De Pasqual et al. (2022), be dependent on the ecological context: we found that Wy females have an advantage under limited male availability, which is similar to the scenario of De Pasqual et al. (2022) in which females were not presented with an alternative male to choose. This suggests that higher male availability may increase intra-sexual competition or affect female behavior and mask Wy females’ reproductive advantage.

Complex life-history trade-offs, such as those between efficient warning color, attractiveness and reproductive success, might be enough to maintain polymorphism in a population (Mérot et al. 2020). Our results highlight the role of sexual selection in maintaining intrapopulation phenotypic variation, by presenting evidence that the male color morphs of the wood tiger moth experience relatively stable differences in their mating success. These mating advantages contrast with a survival advantage in alternative morphs, suggesting that trade-offs between natural and sexual selection may be key to maintaining variation in this species. Although we found little variation in sexual selection between our experimental set-ups, more research on the effects of varying environmental contexts on sexual selection are necessary to fully understand how it may contribute to maintaining polymorphism and affect morph frequencies under changing conditions.

## Acknowledgments

We are extremely thankful to Kaisa Suisto, Jimi Kirvesoja, Teemu Tuomaala, and the greenhouse staff for rearing the moths used in these experiments. We also thank Cristina Ottocento and Linda Hernández Duran for helping with the small cage experiment.

