## Supplemental Figure S1 and Tables S1-S22 for "Exploring mate choice and male strategies in a polymorphic moth"

### SUPPLEMENTARY MATERIAL

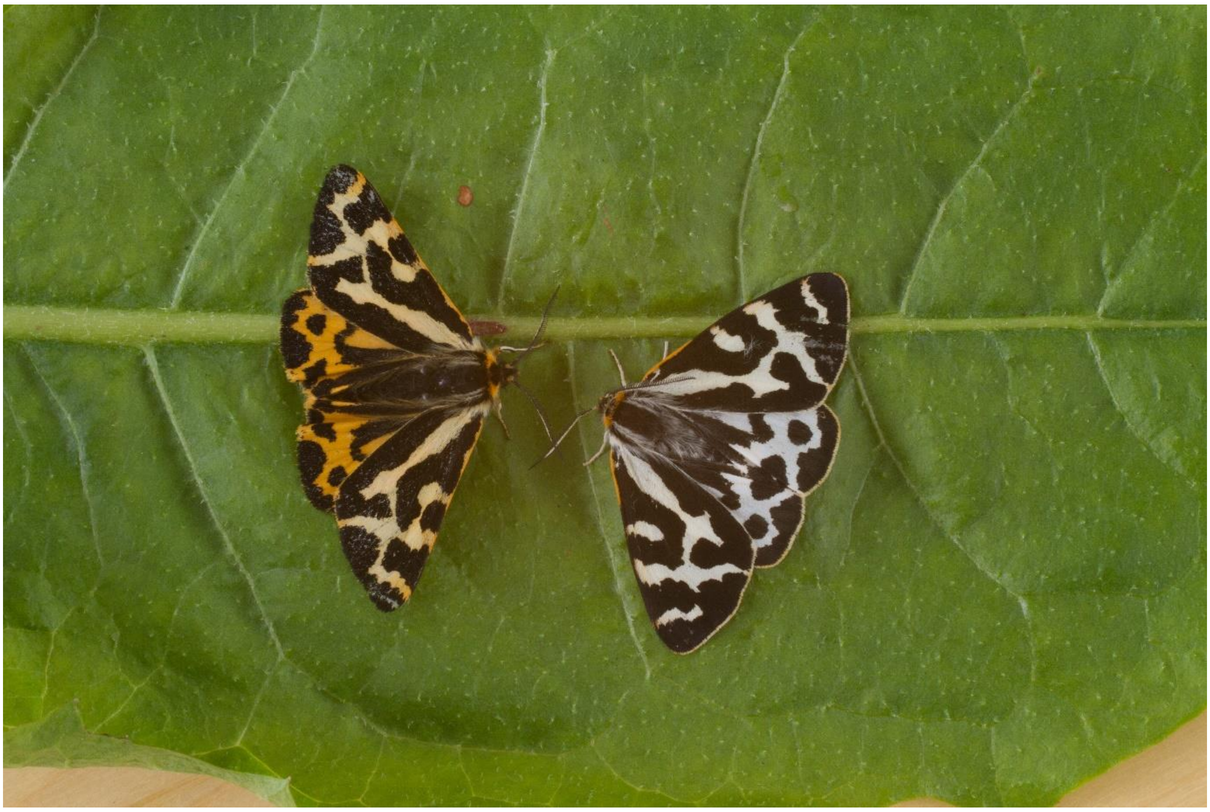

**Figure S1.** A phenotypically yellow (yy) and white (WW or Wy) wood tiger moth male. Image by Samuel Waldron.

**Table S1.** Model selection via AIC for mating success in the pairwise WW – Wy treatment. All models include female ID as a random effect.

| Fixed effects | AIC | AICc |
|---|---|---|
| MaleGenotype + FemaleGenotype + MaleWeight + Generation + MaleGenotype:FemaleGenotype + MaleGenotype:MaleWeight | 114.44 | 115.35 |
| MaleGenotype + FemaleGenotype + MaleWeight + Generation + MaleGenotype:MaleWeight | 111.05 | 111.70 |
| MaleGenotype + FemaleGenotype + MaleWeight + Generation | 109.08 | 109.51 |
| MaleGenotype + MaleWeight + Generation | 108.53 | 108.78 |
| **MaleGenotype + MaleWeight** | **106.65** | **106.78** |
| MaleGenotype | 109.35 | 109.39 |

**Table S2.** Model selection via AIC for mating success in the pairwise WW – yy treatment. All models include female ID as a random effect.

| Fixed effects | AIC | AICc |
|---|---|---|
| MaleGenotype + FemaleGenotype + MaleWeight + Generation + MaleGenotype:FemaleGenotype + MaleGenotype:MaleWeight | 124.16 | 124.84 |
| MaleGenotype + FemaleGenotype + MaleWeight + MaleGenotype:FemaleGenotype + MaleGenotype:MaleWeight | 122.19 | 122.67 |
| MaleGenotype + FemaleGenotype + MaleWeight + MaleGenotype:MaleWeight | 122.25 | 122.57 |
| **MaleGenotype + MaleWeight + MaleGenotype:MaleWeight** | **121.23** | **121.42** |
| MaleGenotype + MaleWeight | 123.51 | 123.60 |
| MaleGenotype | 124.25 | 124.28 |

**Table S3.** Model selection via AIC for mating success in the pairwise Wy – yy treatment. All models include female ID as a random effect.

| Fixed effects | AIC | AICc |
|---|---|---|
| MaleGenotype + FemaleGenotype + MaleWeight + Generation + MaleGenotype:FemaleGenotype + MaleGenotype:MaleWeight | 167.22 | 167.83 |
| MaleGenotype + FemaleGenotype + MaleWeight + Generation + MaleGenotype:MaleWeight | 164.32 | 164.75 |
| MaleGenotype + FemaleGenotype + MaleWeight + Generation | 162.56 | 162.85 |
| MaleGenotype + MaleWeight + Generation | 161.87 | 162.04 |
| MaleGenotype + Generation | 163.18 | 163.27 |
| **MaleGenotype** | **163.32** | **163.35** |

**Table S4.** Model selection via AIC for mating success in the large cage experiment during the first night. All models include replicate as a random effect.

| Fixed effects | AIC |
|---|---|
| MaleGenotype + MaleWeight + MaleGenotype:MaleWeight | 216.26 |
| MaleGenotype + MaleWeight | 212.58 |
| **MaleGenotype** | **211.98** |

**Table S5.** Model selection via AIC for mating success in the large cage experiment during the second night. All models include replicate as a random effect.

| Fixed effects | AIC |
|---|---|
| MaleGenotype + MaleWeight + MaleGenotype:MaleWeight | 131.37 |
| MaleGenotype + MaleWeight | 133.74 |
| **MaleGenotype** | **132.37** |

**Table S6.** Model selection via AIC for likelihood of males being rejected in the pairwise experiment. All models include female ID as a random effect.

| Fixed effects | AIC |
|---|---|
| MaleGenotype + FemaleGenotype + MaleGenotype:FemaleGenotype | 153.33 |
| MaleGenotype + FemaleGenotype | 147.73 |
| **MaleGenotype** | **145.56** |

**Table S7.** Model selection via AIC for number of rejections in the large cage experiment. All models include replicate as a random effect.

| Fixed effects | AIC |
|---|---|
| MaleGenotype + FemaleGenotype + MaleGenotype:FemaleGenotype | 544.66 |
| MaleGenotype + FemaleGenotype | 541.20 |
| **MaleGenotype** | **540.23** |

**Table S8.** Model selection via AIC for mating latency in the pairwise experiment.

| Fixed effects | AIC |
|---|---|
| MaleGenotype + FemaleGenotype + Generation + MaleGenotype:FemaleGenotype | 1049.6 |
| MaleGenotype + FemaleGenotype + Generation | 1043.6 |
| MaleGenotype + Generation | 1040.4 |
| **Generation** | **1037.3** |

**Table S9.** Model selection via AIC for mating latency in the large cage experiment. All models include replicate as random effect.

| Fixed effects | AIC |
|---|---|
| MaleGenotype + FemaleGenotype + Night + MaleGenotype:FemaleGenotype | 639.32 |
| MaleGenotype + FemaleGenotype + Night | 636.42 |
| MaleGenotype + Night | 634.47 |
| **MaleGenotype** | **634.22** |

**Table S10.** Model selection via AIC for viability of offspring in the pairwise experiment.

| Fixed effects | AIC | AICc |
|---|---|---|
| FemaleGenotype + MaleGenotype + FemaleWeight + Generation + FemaleGenotype:MaleGenotype + FemaleGenotype:FemaleWeight | 85.39 | 86.51 |
| FemaleGenotype + MaleGenotype + FemaleWeight + Generation + FemaleGenotype:MaleGenotype | 82.52 | 83.31 |
| FemaleGenotype + MaleGenotype + FemaleWeight + Generation | 80.22 | 80.74 |
| FemaleGenotype + MaleGenotype + Generation | 78.23 | 78.54 |
| **FemaleGenotype + Generation** | **78.36** | **78.51** |
| FemaleGenotype | 90.43 | 90.48 |

**Table S11.** Model selection via AIC for number of larvae in the pairwise experiment.

| Fixed effects | AIC | AICc |
|---|---|---|
| FemaleGenotype + MaleGenotype + FemaleWeight + Generation + FemaleGenotype:MaleGenotype + FemaleGenotype:FemaleWeight | 669.45 | 671.07 |
| FemaleGenotype + MaleGenotype + FemaleWeight + Generation + FemaleGenotype:FemaleWeight | 663.97 | 665.10 |
| FemaleGenotype + FemaleWeight + Generation + FemaleGenotype:FemaleWeight | 660.30 | 661.04 |
| FemaleGenotype + FemaleWeight + Generation | 658.39 | 658.83 |
| **FemaleGenotype + FemaleWeight** | **657.37** | **657.58** |
| FemaleGenotype | 667.95 | 668.02 |

**Table S12.** Model selection via AIC for number of eggs in the pairwise experiment.

| Fixed effects | AIC | AICc |
|---|---|---|
| FemaleGenotype + MaleGenotype + FemaleWeight + Generation + FemaleGenotype:MaleGenotype + FemaleGenotype:FemaleWeight | 930.45 | 931.63 |

| | | |
|---|---|---|
| FemaleGenotype + MaleGenotype + FemaleWeight + Generation + FemaleGenotype:FemaleWeight | 924.47 | 925.30 |
| FemaleGenotype + FemaleWeight + Generation + FemaleGenotype:FemaleWeight | 921.36 | 921.91 |
| **FemaleGenotype + FemaleWeight + FemaleGenotype:FemaleWeight** | **921.21** | **921.53** |
| FemaleGenotype + FemaleWeight | 927.50 | 927.66 |
| FemaleGenotype | 959.01 | 959.06 |

**Table S13.** Model selection via AIC for hatching success in the pairwise experiment.

| Fixed effects | AIC | AICc |
|---|---|---|
| FemaleGenotype + MaleGenotype + FemaleWeight + Generation + FemaleGenotype:MaleGenotype + FemaleGenotype:FemaleWeight | 6.01 | 7.63 |
| FemaleGenotype + MaleGenotype + FemaleWeight + Generation + FemaleGenotype:FemaleWeight | 0.49 | 1.62 |
| FemaleGenotype + FemaleWeight + Generation + FemaleGenotype:FemaleWeight | -2.31 | -1.57 |
| FemaleGenotype + FemaleWeight + Generation | -5.02 | -4.58 |
| FemaleGenotype + Generation | -7.01 | -6.72 |
| **FemaleGenotype** | **-7.88** | **-7.81** |

**Table S14.** Model selection via AIC for viability of offspring in the large cage experiment. All models include replicate as random effect.

| Fixed effects | AIC | AICc |
|---|---|---|
| FemaleGenotype + MaleGenotype + FemaleWeight + Night + FemaleGenotype:FemaleWeight | 68.03 | 69.36 |
| FemaleGenotype + MaleGenotype + FemaleWeight + Night | 66.10 | 66.97 |
| FemaleGenotype + FemaleWeight + Night | 64.31 | 64.82 |
| FemaleGenotype + Night | 62.57 | 62.82 |
| **FemaleGenotype** | **62.94** | **63.02** |

**Table S15.** Model selection via AIC for number of larvae in the large cage experiment. All models include replicate as random effect.

| Fixed effects | AIC | AICc |
|---|---|---|

| | | |
|---|---|---|
| FemaleGenotype + MaleGenotype + FemaleWeight + Night + FemaleGenotype:FemaleWeight | 429.60 | 431.42 |
| FemaleGenotype + MaleGenotype + FemaleWeight + Night | 426.01 | 427.19 |
| FemaleGenotype + FemaleWeight + Night | 422.70 | 423.39 |
| FemaleGenotype + FemaleWeight | 420.99 | 421.32 |
| **FemaleGenotype** | **421.43** | **421.53** |

**Table S16.** Model selection via AIC for number of eggs in the large cage experiment. All models include replicate as random effect.

| Fixed effects | AIC | AICc |
|---|---|---|
| FemaleGenotype + MaleGenotype + FemaleWeight + Night + FemaleGenotype:FemaleWeight | 565.91 | 567.24 |
| **FemaleGenotype + FemaleWeight + Night + FemaleGenotype:FemaleWeight** | **566.94** | **567.81** |
| FemaleGenotype + FemaleWeight + FemaleGenotype:FemaleWeight | 570.84 | 571.35 |
| FemaleGenotype + FemaleWeight | 571.87 | 572.12 |
| FemaleGenotype | 588.37 | 588.45 |

**Table S17.** Model selection via AIC for hatching success in the large cage experiment. All models include replicate as random effect.

| Fixed effects | AIC | AICc |
|---|---|---|
| FemaleGenotype + MaleGenotype + FemaleWeight + Night + FemaleGenotype:FemaleWeight | 1.63 | 3.44 |
| FemaleGenotype + FemaleWeight + Night + FemaleGenotype:FemaleWeight | -2.34 | -1.16 |
| FemaleGenotype + FemaleWeight + Night | -5.74 | -5.05 |
| FemaleGenotype + Night | -7.74 | -7.41 |
| **FemaleGenotype** | **-9.50** | **-9.39** |

**Table S18.** Variance inflation factor (VIF) of all terms included in the model for mating success in the WW-Wy treatment of the pairwise experiment. VIF above 5 is considered moderate collinearity and a VIF above 10 is considered strong collinearity.

| Term | VIF |
|---|---|

| | |
|---|---|
| Male genotype | 1.01 |
| Male weight | 1.01 |

**Table S19.** VIF of all terms included in the model for mating success in the WW-yy treatment of the pairwise experiment.

| Term | VIF |
|---|---|
| Male genotype | 1.25 |
| Male weight | 1.49 |
| Male genotype:Male weight | 1.65 |

**Table S20.** VIF of all terms included in the model for number of larvae in the pairwise experiment.

| Term | VIF |
|---|---|
| Female genotype | 1.02 |
| Female weight | 1.02 |

**Table S21.** VIF of all terms included in the model for number of eggs in the pairwise experiment.

| Term | VIF |
|---|---|
| Female genotype | 1.07 |
| Female weight | 1.83 |
| Female genotype:Female weight | 1.88 |

**Table S22.** VIF of all terms included in the model for number of eggs in the large cage experiment.

| Term | VIF |
|---|---|
| Female genotype | 1.80 |
| Female weight | 5.28 |
| Night | 1.10 |
| Female genotype:Female weight | 6.55 |
